# FLAMs: A self-replicating *ex vivo* model of alveolar macrophages for functional genetic studies

**DOI:** 10.1101/2021.12.12.472259

**Authors:** Sean Thomas, Kathryn Wierenga, James Pestka, Andrew J Olive

## Abstract

Alveolar macrophages (AMs) are tissue resident cells in the lungs derived from the fetal liver that maintain lung homeostasis and respond to inhaled stimuli. While the importance of AMs is undisputed, they remain refractory to standard experimental approaches and high-throughput functional genetics as they are challenging to isolate and rapidly lose AM properties in standard culture. This limitation hinders our understanding of key regulatory mechanisms that control AM maintenance and function. Here, we describe the development of a new model, fetal liver-derived alveolar-like macrophages (FLAMs), which maintains cellular morphologies, expression profiles, and functional mechanisms similar to murine AMs. FLAMs combine treatment with two key cytokines for AM maintenance, GM-CSF and TGFβ. We leveraged the long-term stability of FLAMs to develop functional genetic tools using CRISPR-Cas9-mediated gene editing. Targeted editing confirmed the role of AM-specific gene *Marco* and the IL-1 receptor *Il1r1* in modulating the AM response to crystalline silica. Furthermore, a genome-wide knockout library using FLAMs identified novel genes required for surface expression of the AM marker Siglec-F, most notably those related to the peroxisome. Taken together, our results suggest that FLAMs are a stable, self-replicating model of AM function that enables previously impossible global genetic approaches to define the underlying mechanisms of AM maintenance and function.

## INTRODUCTION

Tissue resident immune cells regulate homeostasis and control local inflammation to external stimuli. A subset of these immune cells are tissue resident macrophages (TRMs) that sample the environment and initiate host responses (1). Distinct TRM populations exist in specific tissues including the liver (Kupffer cells), the skin (Langerhans cells), the brain (microglia), and the lungs (alveolar macrophages [AMs]). These distinct TRMs all have unique functions that are regulated by the local environment and are required for tissue maintenance (2, 3).

As the first line of defense in the airways, AMs are particularly important for tuning the host immune response in the lungs (4). AMs can be distinguished from other macrophage populations in the lung by the surface expression of the sialic acid receptor Siglec-F, the scavenger receptor MARCO, and the integrin CD11c in addition to the high expression and activity of the transcription factor PPARγ, which drives many AM-specific genes (5, 6). AMs are a long-lived and self-replicating and, like most TRMs, are derived from embryonic precursors (7). AMs arise from fetal liver monocytes, which migrate to the lung and develop into mature AMs in the presence of cytokines such as GM-CSF and TGFβ shortly after birth (8–10). The continued presence of these factors is necessary for the maintenance and self-renewal of AMs in the lung, in part by promoting expression and activation of PPARγ (10, 11). Genes and pathways induced by this receptor are involved in lipid metabolism and induction of scavenger receptors that promote phagocytosis (12). This is critical for the AM roles of maintaining surfactant homeostasis, efferocytosis of cellular debris, and phagocytosis of inhaled microbes and particles in the alveolar space (12, 13). Impaired clearance of surfactant by AMs can result in the pathophysiological condition known as pulmonary alveolar proteinosis (PAP) (14). In addition, reduced AM efferocytosis and phagocytosis has been observed in patients with asthma, COPD, and cystic fibrosis, likely contributing to the sustained inflammation and susceptibility to infection observed in these diseases (15–19).

Despite the paramount importance of AMs for lung health, there continue to be key gaps in our understanding of how they are maintained and function to regulate the host response in the lungs. One hurdle towards a mechanistic understanding of AMs is that experiments employing primary AMs require large numbers of animals to isolate a small number of cells that do not robustly proliferate or maintain AM-like functions *ex vivo* (20). This limitation has prevented genetic approaches from being employed to better understand AM maintenance and function. As a result, many *ex vivo* studies investigating responses to airborne particles and microbes rely on bone-marrow derived macrophages (BMDMs) or transformed macrophage cell lines as surrogates of AM biology (21–23). While these macrophage models are useful, they do not faithfully recapitulate all AM functions (24–26). A recent alternative approach cultured cells from the murine fetal liver in the presence of GM-CSF to generate AM-like cells that are functionally and phenotypically like AMs (25). This approach enabled the isolation of large numbers of AM-like cells that might be amenable to tractable genetic approaches. However, we found here that fetal liver-derived cells cultured in GM-CSF alone lost their AM-like morphology, phenotype, and surface marker expression over time, suggesting that GM-CSF is insufficient to maintain the AM-like phenotype. A recent study found that AMs could be continuously cultured *ex vivo* in the presence of GM-CSF and TGFβ (27), which is consistent with reports that TGFβ promotes AM development and maintains AM function both *in vivo* and *ex vivo* (10).

Here, we found that growing fetal liver cells in both GM-CSF and TGFβ results in a long-term stable population of cells that are phenotypically and functionally similar to AMs. Using these fetal liver-derived alveolar-like macrophages (FLAMs), we developed targeted and global genetic tools to dissect regulatory networks that are required to maintain AM-like cells and function. Employing targeted gene-editing, we show here that directed mutations are readily introduced in FLAMs to query specific AM functions. We further demonstrate the utility of FLAMs by using genome-wide CRISPR-Cas9 knockout screen to identify genes that are required for the surface expression of the AM-specific marker Siglec-F. The screen identified key pathways used to maintain Siglec-F expression and the AM-like state including the observation that peroxisome biogenesis plays a central role in maintaining AM functions. Together our results show that FLAMs enable the global dissection of AM regulatory mechanisms at a previously impossible scale.

## MATERIALS AND METHODS

### Animals

Experimental protocols were approved by the Institutional Animal Care and Use Committee at MSU (AUF # PROTO201800113). 6-8 week old C57Bl6 mice (cat # 000664) and Cas9^+^ mice (cat # 026179) were obtained from Jackson Laboratories (Bar Harbor, ME). Mice were given free access to food and water under controlled conditions (humidity: 40–55%; lighting: 12 hour light/dark cycles; and temperature: 24±2°C) as described previously (28, 29). Pregnant dams at 8-10 weeks of age and 14-18 gestational days were euthanized to obtain murine fetuses. AMs were isolated from male and female mice older than 10 weeks of age. BMDMs were obtained from male and female mice 6 weeks of age and older.

### FLAMs cell isolation and culture

Fetal liver derived cells were obtained as previously described (30). Briefly, pregnant dams were euthanized by CO_2_ inhalation for 10 min to ensure death to neonates, which are resistant to anoxia. Cervical dislocation was used as a secondary form of death for the dam. Fetuses were immediately removed, and loss of maternal blood supply served as a secondary form of death for the fetuses. Cells were cultured in complete Roswell Park Memorial Institute medium (RPMI, Thermo Fisher) containing 10% fetal bovine serum (FBS, R&D Systems), 1% penicillin-streptomycin (P/S, Thermo Fisher), 30 ng/mL recombinant mGM-CSF (Peprotech), and 20 ng/mL recombinant hTGFβ1 (Peprotech) included where indicated. Media was refreshed every 2-3 days. When cells reached 70-90% confluency, they were lifted by incubating for 10 minutes with 37°C phosphate-buffered saline (PBS) containing 10 mM EDTA, followed by gentle scraping. After approximately 1 week, adherent cells adopted a round, AM-like morphology. At this time, stocks were frozen for future use. Thawed stocks were plated in untreated Petri dishes with either GM-CSF or GM-CSF and TGFβ (20 ng/mL recombinant hTGFβ1 [Peprotech]) and sub-cultured as described above.

### AM isolation and culture

Mice were euthanized by CO_2_ exposure followed by exsanguination via the inferior vena cava. Lungs were lavaged as previously described (20). Cells were then resuspended in RPMI media containing 30 ng/mL GM-CSF and plated in untreated 48- or 24- well plates. AMs were lifted from plates using Accutase™ (BioLegend) and seeded for experiments.

### BMDM isolation and culture

C57BL/6J mice were euthanized by CO_2_ exposure followed by cervical dislocation. Both femurs were cut on one end to expose the bone marrow, placed cut side down in 0.6 mL tubes, and centrifuged at 16,000 *x*g for 25 seconds. Marrow from multiple mice was pooled, dissociated to a single cells suspension in sterile PBS and pelleted by centrifuging at 220 *x*g for 5 min. The pellet was resuspended in mouse RBC lysis buffer (Alfa Aeser) and incubated at room temperature for 5 minutes. The RBC lysis buffer was diluted with 2 volumes of PBS and the cell suspension passed through a nylon 70 μm filter (Corning). Cells were pelleted a second time and resuspended in RPMI media containing 10% FBS, 1% P/S, and 20% L929 media (31). Approximately 5 x 10^6^ cells were plated per dish in 10 cm untreated petri dishes. Media was refreshed every 2-3 days. Cells were used for assays when fully differentiated after 7 days.

### Flow cytometry

Plated cells were lifted in warm PBS with 10 mM EDTA for 5-10 minutes and washed twice in PBS before fluorescent antibody labeling. Immediately following isolation, AMs were resuspended in PBS and filtered through a 70 μm basket filter and incubated with an antibody cocktail of PE CD170, APC CD11c, APC-Cyanine7 CD14, and FC Block (Biolegend; 1:400 in PBS) for 20 min at room temperature in light-free condition. Immunochemically labeled cells were washed three times with PBS, resuspended in PBS, and passed through a 70 μm nylon filter immediately prior to analysis. Flow cytometry was performed on a LSR II Flow Cytometer (BD Biosciences) at the Michigan State University Flow Cytometry Core.

### qPCR

RNA was isolated from ~5 x 10^5^ cells using RNeasy mini kits (Qiagen), typically yielding 100-400 ng RNA. RNA was then reverse transcribed to cDNA using a High-Capacity cDNA reverse transcription kit (Thermo Fisher) on a Stratagene Robocycler 40. Quantitative real-time qPCR was performed using specific Taqman probes (Thermo Fisher) for TGFβ1 (*Tgfb1*), TGFβ receptors (*Tgfbr1, Tgrbr1*), selected genes used to distinguish AMs from other macrophage populations (*Cd14, Siglecf, Marco, Pparg Car4, Fabp4, Itgax*), and cytokines (*Il1a, Il1b, Il10*) on an Applied Biosystems™ QuantStudio™ 7 real-time PCR system. Data were analyzed with Applied Biosystems™ Thermo Fisher Cloud using the RQ software and the relative quantification method. *Gapdh* was used as the housekeeping gene. Relative copy number (RCN) for each gene was normalized to expression of *Gapdh* and calculated as described previously (32).

### Scanning electron microscopy

Suspensions of AMs or FLAMs were diluted to 2.5 x 10^5^ cells/mL, and 100 μL pipetted directly upon glass 12 mm diameter, 0.13-0.16 mm thick circular coverslips (Electron Microscopy Sciences), which were placed in the bottom of 6-well plates. Cells were allowed to settle for 2-3 minutes, then 1 mL of media was added to fill the well. To fix cells, the coverslips were removed from the wells, submerged in 4% glutaraldehyde in 0.1 M sodium phosphate buffer at pH 7.4 and placed in a graded ethanol series (25%, 50%, 75%, 95%) for 10 min at each step followed by 3 minutes changes in 100% ethanol.

Samples were critical point dried in a Leica Microsystems model EM CPD300 critical point drier (Leica Microsystems, Vienna, Austria) using CO_2_ as the transitional fluid. Coverslips were then mounted on aluminum stubs using epoxy glue (System’s Three Quick Cure 5, Systems Three Resins, Auburn WA). Samples were coated with osmium at ~10 nm thickness in an NEOC-AT osmium chemical vapor deposition coater (Meiwafosis Co, Osaka, Japan) and examined in a JEOL 7500F (field emission emitter) scanning electron microscope (JEOL, Tokyo, Japan).

### cSiO_2_ phagocytosis assay

To assess phagocytosis of cSiO_2_ particles, FLAMs, AMs, and BMDMs were seeded at 0.25 cells/cm^2^ in 48- or 96-well plates to observe engulfment of surrounding silica particles. The following day, the media was removed, wells were rinsed 1x with sterile PBS, media replaced with FluoroBrite DMEM (Thermo Fisher) containing 10% FBS and 200 nM SYTOX Green nucleic acid stain (Thermo Fisher). cSiO_2_ was then added dropwise to a final density of 25-100 μg/cm_2_. Cells were imaged over time on an EVOS FL2 fluorescent microscope (Thermo Fisher) with an on-stage, temperature control CO_2_ incubator and 2-4 images were acquired per well. SYTOX Green detected on the GFP light cube.

Images were analyzed using analysis pipelines built in the CellProfiler software (33). cSiO_2_ engulfment was assessed by quantifying the number of cSiO_2_-filled cells, which have a higher pixel intensity than non-cSiO_2_-filled cells due to the accumulation of the particles. To avoid counting aggregated cSiO_2_ particles, a threshold was applied to capture only shapes with high solidity and low compactness. Cell death was quantified by counting SYTOX Green^+^ cells, respectively.

### ELISAs

Cells were treated with cSiO_2_ for 8 hours or LPS for 24 hours at the indicated concentrations. Cell-free supernatant was collected and the cytokines IL-1α, IL-1β, and IL-10 were analyzed using DuoSet ELISA kits (R&D Systems) per the manufacturer’s instructions.

### CRISPR Targeted Knockouts

sgRNA cloning sgOpti was a gift from Eric Lander & David Sabatini (Addgene plasmid #85681) (34). Individual sgRNAs were cloned as previously described (35). In short, sgRNA targeting sequences were annealed and phosphorylated then cloned into a dephosphorylated and BsmBI (New England Biolabs) digested SgOpti. sgRNA constructs were then packaged into lentivirus as previously described and used to transduce early passage FLAMs. Two days later, transductants were selected with puromycin. After one week of selection, gDNA was isolated from each targeted FLAM, and PCR was used to amplify edited regions and sanger sequencing was used to quantify indels.

### Construction of genome-wide loss-of-function library and Siglec-F Screen

The mouse BRIE knockout CRISPR pooled library was a gift of David Root and John Doench (Addgene #73633) (36). Using the BRIE library, 4 sgRNAs targeting every coding gene in mice in addition to 1000 non-targeting controls (78,637 sgRNAs total) were packaged into lentivirus using HEK293T cells and transduced Cas9^+^ FLAMs at a low multiplicity of infection (MOI <0.3). Two days later these cells were selected with puromycin. We then passaged the transduced library in TGFβ in parallel with non-transduced cells of the same passage without TGFβ. When the non-transduced cells grown in the absence of TGF–β showed reduced Siglec-F expression by flow cytometry we isolated gDNA from the library for sequencing and found high coverage and distribution (**Table S1**). In parallel, the transduced library was fixed, and fluorescence activated cell sorting (FACS) was used to isolate the SiglecF^high^ and SiglecF^low^ bins using a BioRad S3e cell sorter. Genomic DNA was isolated from each sorted population from two biological replicate experiments using a homemade modified salt precipitation method previously described (37). Amplification of sgRNAs by PCR was performed as previously described using Illumina compatible primers from IDT (36), and amplicons were sequenced on an Illumina NovaSeq 6000 at the RTSF Genomics Core at Michigan State University.

Sequence reads were first trimmed to remove any adapter sequence and to adjust for p5 primer stagger. We used MAGeCK to map reads to the sgRNA library index without allowing for any mismatch. Subsequent sgRNA counts were median normalized to control sgRNAs in MAGeCK to account for variable sequencing depth. To test for sgRNA and gene enrichment, we used the ‘test’ command in MAGeCK to compare the distribution of sgRNAs in the SiglecF^high^ and SiglecF^low^ bins.

### Bioinformatic analysis

Both DAVID analysis and GSEA analysis were used to identify enriched pathways and protein families that were enriched in the data set. Genes were ranked in MAGeCK using RRA and the top enriched positive regulators (4-fold change with at least 2 sgRNAs) were used as a “candidate list” in both DAVID analysis using default settings (38). Functional analysis and functional annotation analysis were completed, and top enriched pathways and protein families were identified. For GSEA analysis, the “GSEA Preranked” function was used to complete functional enrichment using default settings for KEGG, Reactome and GO terms.

### Data availability

Raw sequencing data in FASTQ and processed formats will be available for download from NCBI Gene Expression Omnibus (GEO) and available upon request.

### Statistical analysis and data visualization

Statistical analysis and data visualization were performed using Prism Version 8 (GraphPad) or R studio as indicated in the figure legends. SYTOX^+^ and cSiO_2_-filled cells were quantified using CellProfiler. Data are presented, unless otherwise indicated, as the mean ± the standard deviation. For parametric data, one-way ANOVA followed by Tukey’s post-hoc test was used to identify significant differences between multiple groups, and Student’s t-tests were used to compare two groups. Non-parametric one-way ANOVAs and Mann-Whitney U tests were used to compare multiple groups and two groups, respectively, for non-parametric data.

## RESULTS

### Fetal liver-derived cells and AMs cultured in GM-CSF alone do not stably express lineage-specific markers over time

The development of a genetically tractable AM *ex vivo* model requires a stable population that maintains AM-like phenotypes and functions long-term. As a first step, we examined the long-term stability of AM-like cells using a previously described method culturing fetal liver derived cells in the cytokine GM-CSF (25). Consistent with previous reports, we found that fetal liver cells grown in GM-CSF phenotypically and morphologically resemble AMs (25, 39, 40). Fetal liver cells grown for 2 weeks *ex vivo* in the presence of GM-CSF adopt a distinct fried-egg-like morphology akin to AMs (**Figure 1A**). Scanning electron microscopy revealed that the surfaces of both AM and low passage fetal liver cells (<1 month of culture) have numerous outer membrane ruffles (**Figure 1B**). However, high passage fetal liver cells (>1 month of culture) underwent a morphological shift from an AM-like, ovoid morphology with numerous outer plasma membrane ruffles, to a smaller, fusiform morphology with loss of membrane ruffles (**Figure 1A and 1B**). Thus, in our hands, the morphology of fetal liver derived cells grown in recombinant GM-CSF are not stable long-term.

**Figure 1.**
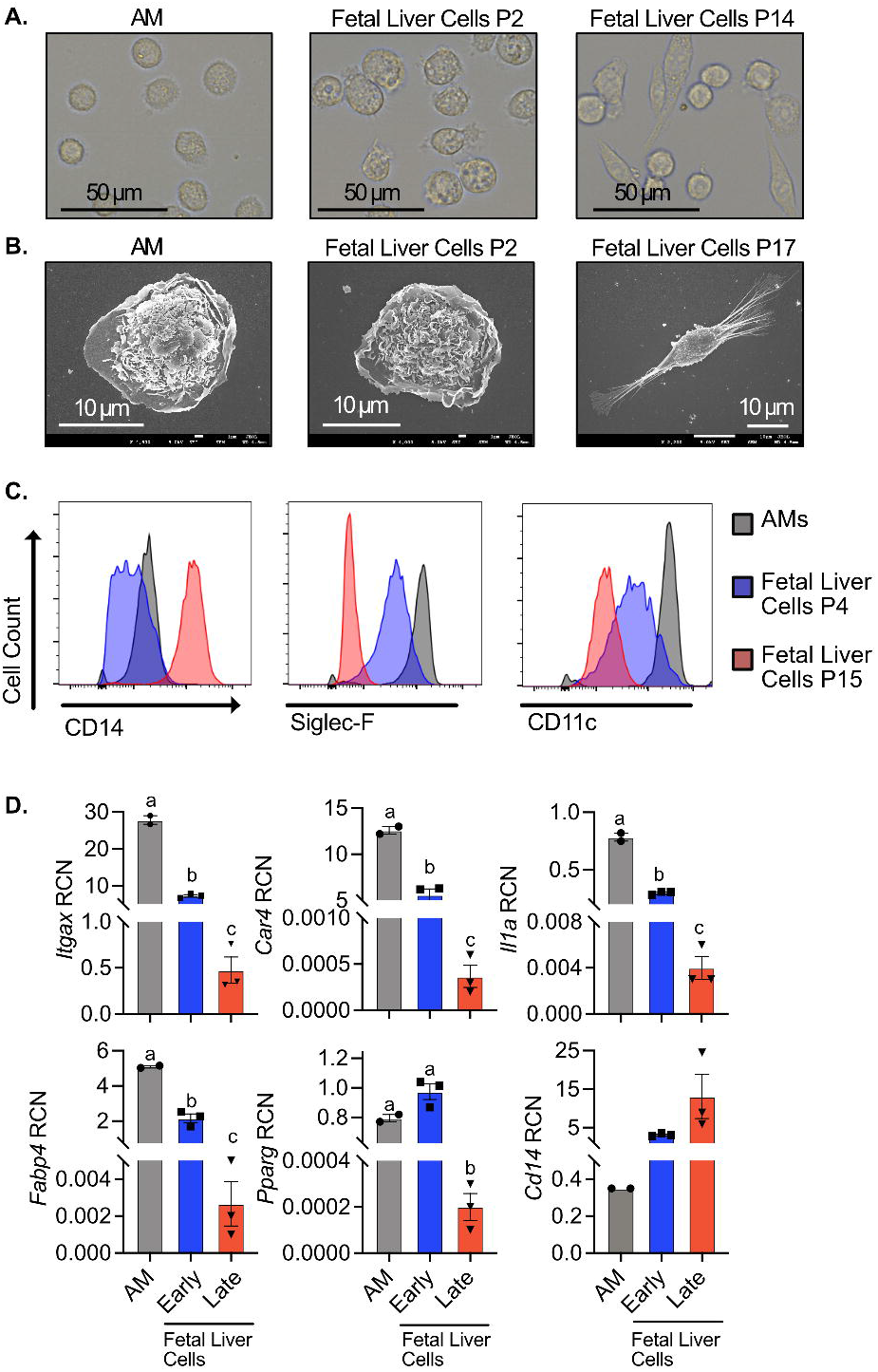
Fetal liver macrophages cultured with GM-CSF lose their AM-like phenotype over time. Fetal liver cells were cultured with GM-CSF and analyzed at indicated passage. AMs were isolated and analyzed immediately. **A)** AMs, P2 fetal liver cells, and P37 fetal liver cells were lifted from culture and imaged on a EVOS FL Auto 2 fluorescent microscope at 60x magnification. **B)** AMs, P3 fetal liver cells, and P15 fetal liver cells were fixed and imaged by scanning electron microscopy at 4500x, 4000x, and 2200x, respectively. **C)** Alveolar macrophages (Gray), P4 fetal liver macrophages (Blue), and P15 fetal liver macrophages (Red) were assessed for surface expression of the markers CD14, Siglec-F, and CD11c by flow cytometry. **D)** Gene expression of indicated genes in AMs, early fetal liver macrophages (P1), and late fetal liver macrophages (P18) was quantified by qPCR. Data was compared using one-way ANOVA followed by Tukey’s multiple comparisons test. Bars labeled with unique letters are significantly different (*p*<0.05). Results are representative of 2 or 3 independent experiments.

We further examined whether changes in surface markers or gene expression varied as fetal liver cells were cultured over time. Using flow cytometry, we found similarity between AMs and low passage fetal liver cells with high surface expression of Siglec-F and CD11c and low expression of CD14. However, high passage fetal liver cells showed low expression of SiglecF and CD11c, while expressing high levels of CD14 (**Figure 1C**). Similarly, when we quantified gene expression, we observed that low passage fetal liver cells and AMs express high levels of *Pparg, Car4, Il1a* and *Fabp4* (a transcriptional target of PPARγ) and low levels of CD14, while high passage fetal liver cells expressed very low levels of the AM-associated transcripts but high levels of CD14 (**Figure 1D**). We observed similar results with AMs isolated from the lungs (**Figure S1A and B**). To exclude the possibility of contaminating cells outcompeting the alveolar-like cells over long-term culture we used fluorescence activated cell sorting (FACS) to isolate a pure Siglec-F positive population of cells that were then cultured in GM-CSF media. We continued to observe a decline in Siglec-F and CD11c in these cells (**Figure S2**). Together these data suggest that prolonged culture of fetal liver-derived cells in GM-CSF media results in a decline in AM-like properties.

### Fetal liver-derived cells grown in GM-CSF and TGFβ are phenotypically similar to AMs long-term

We next pursued strategies to improve the stability of AM-specific phenotypes of fetal liver-derived cells grown *ex vivo*. Based on a prior report that the cytokine TGFβ is critical to AM development and homeostasis (10), we hypothesized that the addition of TGFβ to our culture system would maintain cells in an AM-like state. We first tested whether the addition of TFGβ alters the expression of AM-associated genes. Fetal liver cells in GM-CSF media were treated for 24 hours with 10 ng/mL TGFβ which we found induced the genes *Pparg, Car4* a transcriptional target of PPAR-γ, and *Itgax* (**Figure S3A**), all of which are highly expressed by AMs (**Figure 1D**). We next examined if continued supplementation with TGFβ stabilizes the AM-like phenotypes of fetal liver cells long-term. Fetal liver-derived cells were cultured in GM-CSF media or GM-CSF media containing 20 ng/mL TGFβ. After 15 passages (approximately 2 months of culturing), cells grown in the presence of TGFβ retained a round, AM-like morphology (**Figure 2A**) and continued expressing AM-identifying genes (**Figure 2B**). Conversely, fetal liver-derived cells cultured without TGFβ lost the expression of AM-identifying genes *Siglecf, Marco* and *Pparg* and began expressing *Cd14*, which is a common marker for monocyte-derived macrophages recruited to the lung (41) (**Figure 2B**).

**Figure 2.**
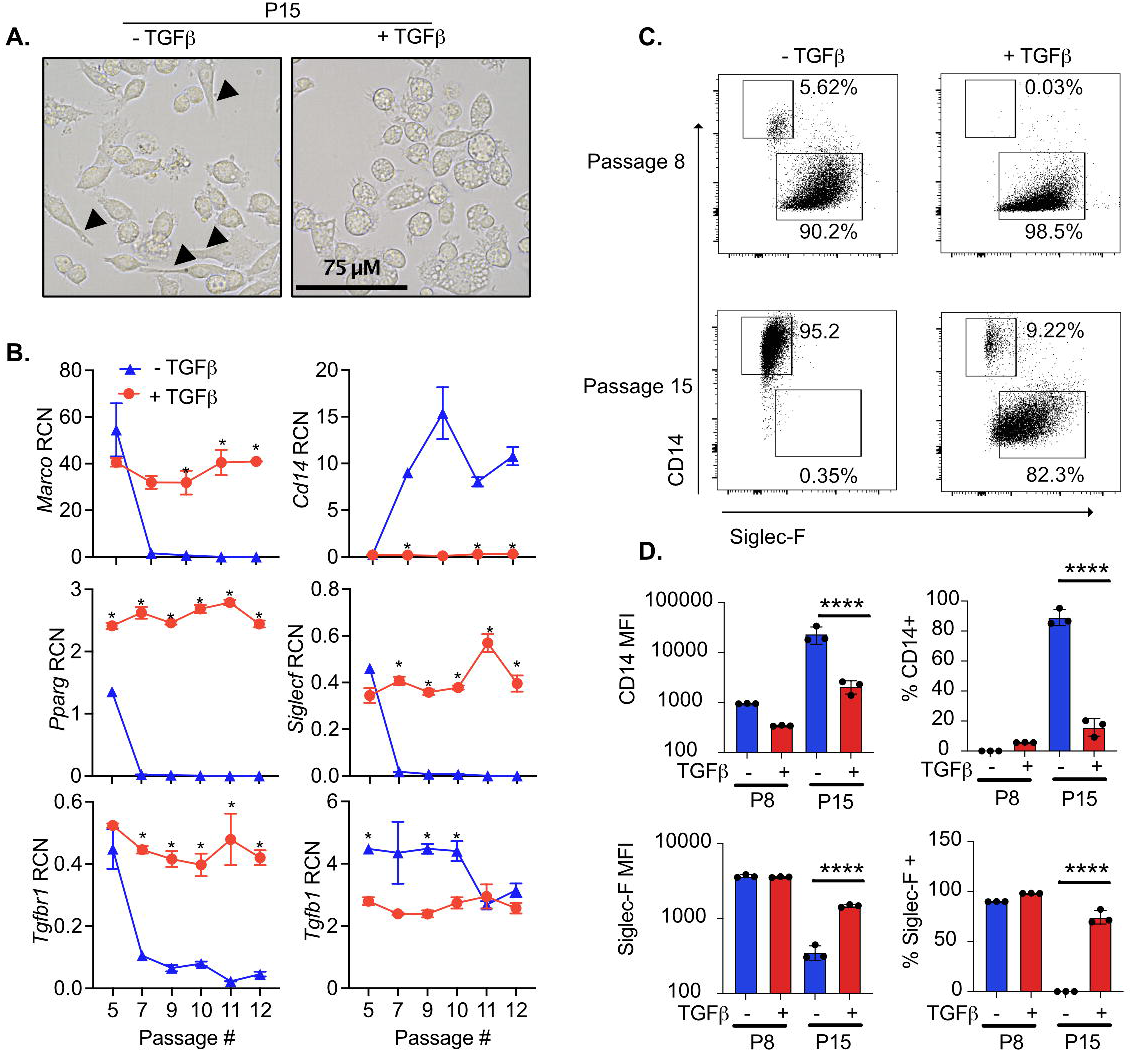
Culturing fetal liver cells with TGFβ and GM-CSF maintains AM-like phenotypes long-term. Fetal liver cells were cultured with or without TGFβ for the indicated passages. **A)** P15 cells cultured with and without TGFβ were imaged on a EVOS FL Auto 2 fluorescent microscope at 60x magnification. Cells with a clearly visible spindleoid morphology are marked with arrows. **B)** At the indicated passage, RNA was extracted from a subset of cells for gene expression analysis. Expression of the indicated genes are quantified as relative copy number (RCN) compared to Gapdh. Asterisks indicate significant (*p*<0.05) differences in gene expression of cells cultured with and without TGF-β cells at the same passage number, as determined by Student’s t-test. **C)** At the indicated passage, cells were analyzed by flow cytometry for the surface expression of CD14 and Siglec-F. Representative biaxial plots from triplicate samples is shown. **D)** Quantification of the mean fluorescence intensity (MFI) and percent of cells positive of CD14 and Siglec-F surface expression from cells in (C) expressed as MFI (left) and percent positive (right). Results are representative of at least two independent experiments. ****p<.0001 by one-way ANOVA with a Tukey correction for multiple comparisons.

We next determined if fetal liver-derived cells grown in the absence of TGFβ would revert to the AM-like state upon the addition of TGFβ. Fetal liver cells grown in the absence of TGFβ were cultured with and without TGFβ for 6 days and the expression of Siglec-F and CD14 was quantified by flow cytometry (**Figure S3B**). In parallel, fetal liver cells maintained in TGFβ were cultured for six days in the presence or absence of TGFβ. We observed that while the removal of TGFβ resulted in a significant decrease in Siglec-F and an increase in CD14, there was no change in expression upon the addition of TGFβ to fetal liver cells that previously lost AM-like marker expression. When we examined the gene expression of *Tgfb1* and the TGFβ receptors, *Tgfbr1* and *Tgfbr2* we observed a significant decrease in expression of the *Tgfb1* (**Figure 2B, Figure S3C**). These data suggest that the loss of AM-like potential of fetal liver cells grown in the absence of TGFβ is not reversible.

We next quantified changes in the surface expression of Siglec-F and CD14 by flow cytometry in the fetal liver-derived cells grown in the presence and absence of TGFβ. Consistent with our gene expression analysis we observed that fetal liver-derived cells lose the expression of Siglec-F and gain the expression of CD14 over time (**Figure 2C and 2D**). In contrast, fetal liver-derived cells grown with TGFβ maintained over 80% of cells with high levels of Siglec-F expression and low levels of CD14. Thus, culturing fetal-liver cells in both GM-CSF and TGFβ results in the stable gene expression of self-replicating cells that phenotypically resemble AMs.

### Fetal liver-derived cells grown in GM-CSF and TGFβ are functionally similar to AMs in response to cSiO_2_ relative to phagocytosis, IL-1 cytokine release, and death

To assess the functional similarity of fetal liver-derived cells grown in TGFβ with AMs, we assessed the response of cells to crystalline silica (cSiO_2_), a respirable particle associated with silicosis and autoimmunity (42, 43). Cells were exposed to various concentrations of cSiO_2_ for 8 hours. SYTOX Green, a membrane impermeable nucleic acid stain, was included to assess lytic cell death. AMs and low passage fetal liver-derived cells grown with and without TGFβ showed similar rates of cSiO_2_ engulfment and cell death (**Figure 3A**). However, late passage fetal liver-derived cells without TGFβ exhibited poor phagocytosis, and as a result, tolerated the presence of cSiO_2_ without inducing cell death. This unresponsiveness is prevented by TGFβ, as late passage fetal liver-derived cells effectively phagocytosed cSiO_2_ (**Figure 3A and 3B**). Rates of phagocytosis by BMDMs were comparable to AMs but were accompanied by a two-fold increase in cell death. Thus, fetal liver-derived cells grown in TGFβ and GM-CSF are functionally stable long-term and recapitulate phagocytosis and cell death kinetics similarly to AMs.

**Figure 3.**
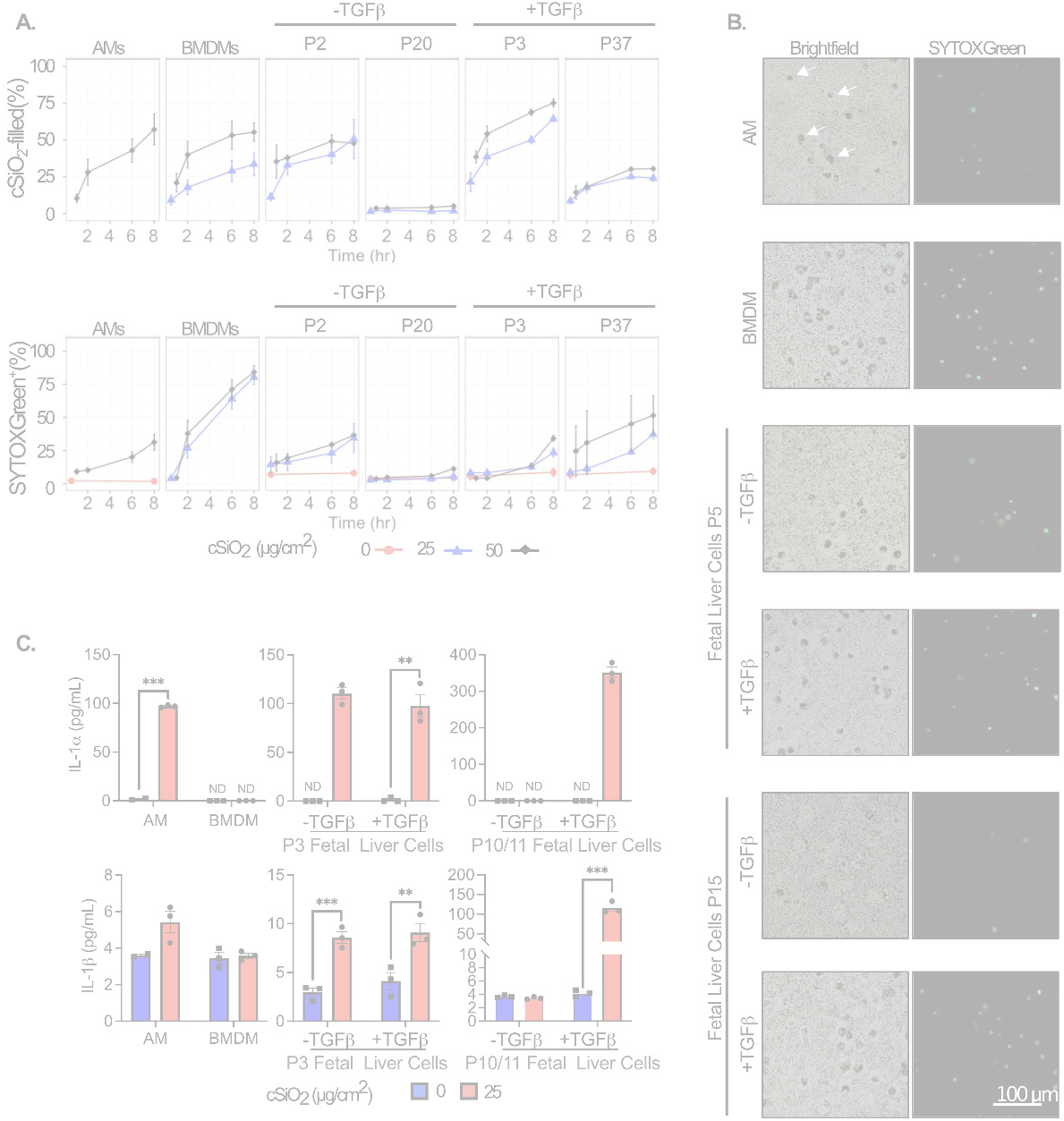
The kinetics of cSiO_2_ uptake and cSiO_2_-induced cell death and IL-1 release are similar among AMs and FLAMs. AMs, bone marrow-derived macrophages (BMDMs), and fetal liver cells were seeded in 96-well plates. After 24 hours, media was replaced with FluoroBrite DMEM containing 200 nM SYTOX green and 10% FBS. cSiO_2_ at the indicated densities was added dropwise to cells and images were taken at 0, 2, 6, and 8 hours using an EVOS FL2 fluorescent microscope. **A)** The percent of cSiO_2_-filled and SYTOX^+^ cells were quantified using the CellProfiler software. **B)** Representative images of SYTOX^+^ and cSiO_2_-filled cells (white arrows in AM panel, top right) treated with 50 μg/cm^2^ silica for 8 hours, 20x magnification. **C)** In a separate experiment, the supernatant was collected after 8 hr treatment with 25 μg/cm^2^ to assess release of the cytokines IL-1α **(Top)** and IL-1β **(Bottom)** by ELISA. Asterisks indicative of significant differences between groups (\*\**p*<0.01, \*\*\**p*<0.001), as assessed by Student’s t-tests between relevant groups. ND = not detected. Results are representative of at least two independent experiments.

IL-1α is associated with the inflammatory response to particle-induced inflammation. *in vivo* and *ex vivo* studies suggest AMs are the primary source of IL-1α in the lung following inhalation of cSiO_2_, likely as a result of cell death (44, 45). Initial characterizations of fetal liver-derived cells grown in GM-CSF (30) showed they respond to LPS like AMs by producing high levels of IL-1α and low levels of IL-10 in contrast BMDMs that make little IL-1α. We replicated these experiments and observed similar results with low passage fetal liver-derived cells producing high levels of IL-1α and low levels of IL-10 in response to LPS (**Figure S3D**). We next tested how the IL-1α response to cSiO_2_ differed over-time in fetal liver-derived cells grown in the presence and absence of TGFβ. We found that high levels of IL-1α were released both low and high passage fetal liver-derived cells grown in both GM-CSF and TGFβ following cSiO_2_ exposure for 8 hours, similar to AMs (**Figure 3C**). In contrast, we observed that cSiO_2_ induced IL-1α release from low passage fetal liver-derived cells grown in GM-CSF alone but not late passage cells. Late passage cells grown in GM-CSF alone instead phenocopied BMDMs and released no detectable IL-1α following cSiO_2_ exposure. Release of IL-1β in these cells may be indicative of inflammasome activation, which is a major mechanism of AM toxicity following exposure to cSiO_2_. We found cSiO_2_ exposure to elicit modest IL-1β release from low passage fetal liver-derived cells grown without TGFβ, and from both low and high passage fetal liver-derived cells grown with TGFβ. We further observed a slight, though not significant, increase in IL-1β release in AMs following cSiO_2_ exposure (**Figure 3C**). cSiO_2_-induced IL-1β release was not evident from BMDMs or late MPI cells. Taken together, these experiments show that growth of fetal-liver cells in both GM-CSF and TGFβ recapitulates many aspects of AM physiology and function as stable, long-term, self-propagating cells. We call these cells <u>F</u>etal <u>L</u>iver-derived <u>A</u>lveolar <u>M</u>acrophages (FLAMs).

### CRISPR-Cas9 editing in FLAMs enables disruption of AM-specific responses to cSiO2

A significant hinderance in the study of AMs is their intractability to standard genetic approaches. This shortcoming has limited the understanding of pathways and regulators that control AM maintenance and function. We hypothesized that FLAMs could be leveraged to dissect AM functional mechanisms. To test this hypothesis we developed CRISPR-Cas9 mediated gene-editing tools by generating FLAMs from Cas9+ mice (46, 47). Using these cells, we targeted *Marco* and *Il1r1*, two genes associated with phagocytosis and inflammatory responses to cSiO_2_ in AMs (48, 49). Each gene was targeted using two independent sgRNAs per gene by lentiviral transduction. Following selection of successfully transduced cells, we evaluated the editing efficiency of each target genes using Tracking of Indels by DEcomposition (TIDE) analysis. We observed robust editing for both sgRNAs with at least one sgRNA per gene reaching over 95% editing efficiency (**Figure S5A,B**). Thus, FLAMs are amenable to genetic targeting by CRISPR-Cas9.

Given the scavenger receptor MARCO has been shown to be involved in cSiO_2_ uptake and toxicity while IL1R1 is known to amplify inflammatory cues, we hypothesized that cells deficient in MARCO and IL1R1 expression would have a reduced inflammatory response to cSiO_2_ (48, 50). We therefore tested whether FLAMs targeted for *Marco* or *Il1r1* would differentially respond to cSiO_2_ exposure compared to wild-type. We exposed control FLAMs and sgMarco or sgIl1r1 FLAMs to two different cSiO_2_ concentrations and quantified cell death. While we observed no change in cell death in sgIl1r1 FLAMs compared to control FLAMs, a significant reduction in cell death in sgMarco FLAMs was observed following high cSiO_2_ exposure (**Figure 4A**). We next examined the production of IL1 following exposure of cells to cSiO_2_. We found reduced cSiO_2_-induced IL-1α and IL-1β production by sgMarco and sgIl1r1 FLAMs compared to control FLAMs (**Figure 4B,C**). Therefore, FLAMs are genetically tractable and can be used to dissect AM-specific functions.

**Figure 4.**
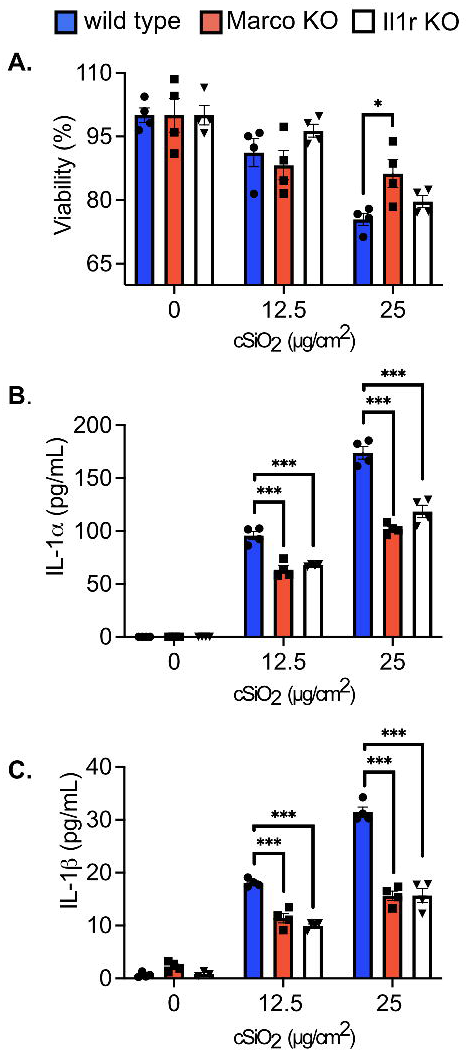
The loss of Marco and Il1r modulate the response of FLAMs to cSiO_2_ treatment. Wild type, *Marco KO*, and *Il1r1 KO* FLAMs were treated with cSiO_2_ at the indicated concentrations for 8 hours. A) Cell viability was determined using the MTS assay, with 100% viability determined as the mean absorbance of the formazan dye product in the untreated WT cells. Supernatant was collected to measure release of B) IL-1α and C) IL-1β. Asterisks indicate significant differences (\**p*<0.05, \*\*\**p*<0.001) between cell types within treatment groups, as determined by one-way ANOVA. Results are representative of at least two independent experiments with biological triplicates.

### Forward genetic screen in FLAMs identifies regulators of the AM surface marker Siglec-F

The genetic tractability of FLAMs opens the possibility of performing forward genetic screens in an AM context, which was previously unviable. We recently developed a screening platform in immortalized bone marrow macrophages (iBMDMs) that uses cell sorting of CRISPR-Cas9 targeted cells to enrich for genes that positively or negatively regulate the surface expression of important immune molecules (51). We hypothesized this screening pipeline could be leveraged to dissect pathways responsible for the unique expression profiles seen in AMs and FLAMs. As a first step to test this hypothesis, we dissected the changes in the surface expression of Siglec-F when targeted using CRISPR-Cas9. Among macrophages, Siglec-F is uniquely expressed on the surface of AMs, yet how Siglec-F is regulated remains entirely unknown. Given that Siglec-F expression is lost as cells lose their AM-like phenotypes, globally understanding Siglec-F regulation in FLAMs may inform key gene networks in AMs. To test the dynamic range of Siglec-F expression on FLAMs, we targeted Siglec-F with two independent sgRNAs in both Cas9^+^ FLAMs and iBMDMs. Again, extensive editing for both sgRNAs was observed with one sgRNA reaching over 99% editing efficiency (**Figure S5C**). As expected, control iBMDMs showed no surface Siglec-F expression and targeting Siglec-F showed no observable change by flow cytometry (**Figure 5A**). In contrast, we observed robust Siglec-F expression on control FLAMs while *sgSiglecF* FLAMs showed a greater than 100-fold reduction in MFI (**Figure 5B**). This dynamic range is comparable to other surface markers we previously screened in iBMDMs, suggesting that Siglec-F is an ideal target for a genetic screen in FLAMs (51).

**Figure 5.**
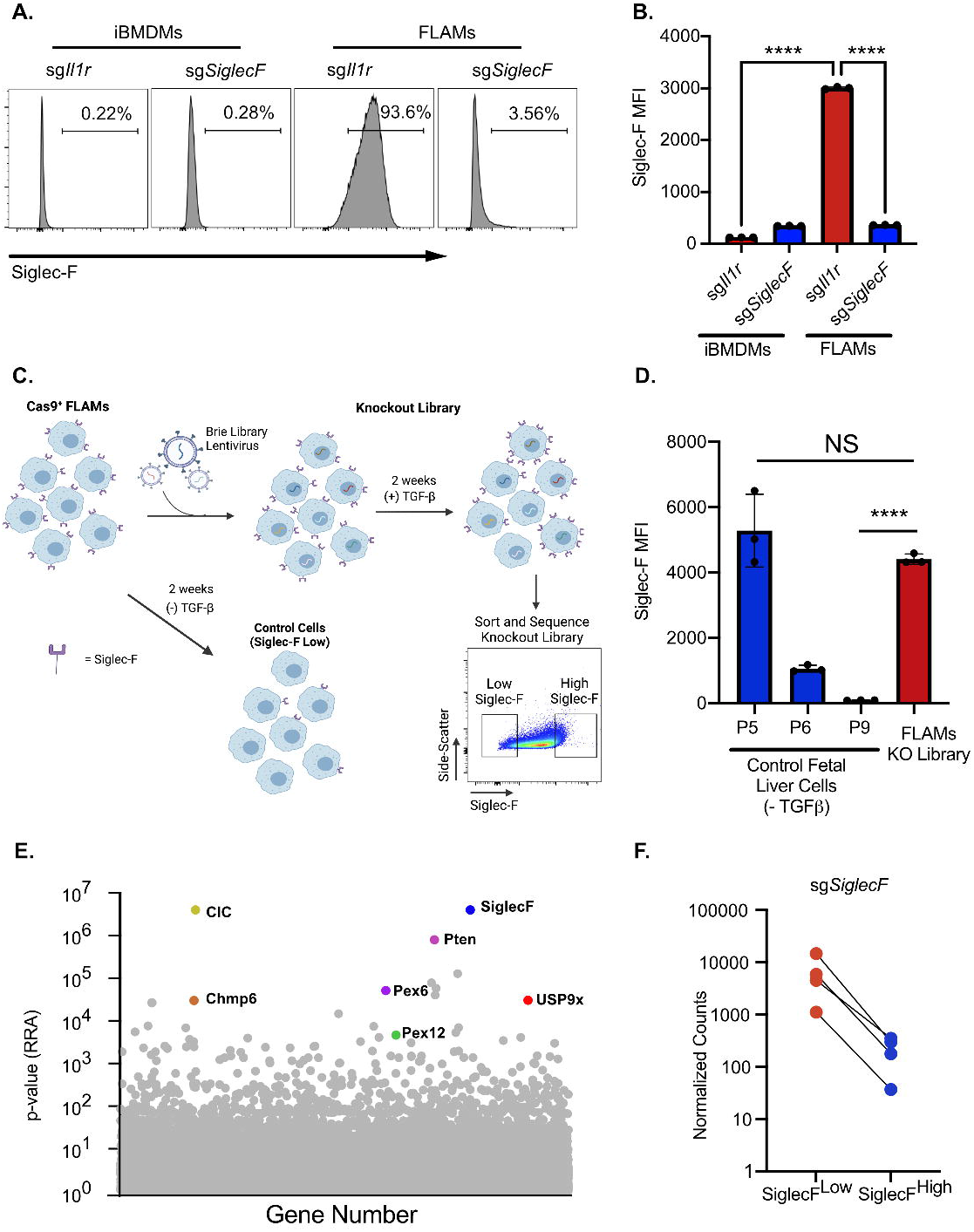
A loss-of-function forward genetic screen identifies regulators of Siglec-F surface expression on FLAMs. iBMDMs or FLAMs targeted for *Il1r1* or *SiglecF* were analyzed by flow cytometry for Siglec-F surface expression. **A)** Shown are representative histograms of surface expression. **B)** The MFI of Siglec-F surface expression was quantified on cells of the indicated genotypes. \*\*\*\**p*<0.0001 between samples by one-way ANOVA with a Tukey correction. These data are representative of two independent experiments **C)** Shown is a schematic of the generation of the FLAM knockout library and screen to identify Siglec-F regulators. The transduction of Cas9+ FLAMs with genome-wide library of sgRNAs results in variable Siglec-F surface expression. When parallel control FLAMs grown in the absence of TGFβ lost Siglec-F expression the top and bottom 5% of Siglec-F expression cells were isolated from the knockout FLAM library by FACS. Sorted cells were then used for downstream sequencing and analysis. **D)** Siglec-F surface expression of library control cells grown in the absence of TGFβ was monitored over time and compared to the transduced FLAM library prior to sorting. Shown is the MFI for Siglec-F expression of the indicated cells and passage numbers **E)** Shown is the α-RRA score of each gene in CRISPR library that passed filtering metrics in MAGeCK. Genes of interest are noted. **F)** Normalized *sgSiglecF* counts for each sgRNA found in both the SiglecF^low^ and SiglecF^high^ sorted populations is shown.

To globally identify genes that contribute to Siglec-F surface expression on FLAMs, we generated a genome-wide knockout library. FLAMs from Cas9^+^ mice were transduced with sgRNAs from the pooled Brie library (52) which contains 4 independent sgRNAs per mouse coding gene. In parallel to the library, we grew control Cas9^+^ fetal liver cells with GM-CSF alone to monitor the loss of Siglec-F expression in the absence of TGFβ signaling (**Figure 5C and 5D**). When control cells lost Siglec-F expression, genomic DNA from the FLAMs knockout library was purified and the sgRNAs were quantified by deep sequencing. The library coverage was confirmed to have minimal skew (**Table S1**). We then conducted a forward genetic screen using FACS to isolate the Siglec-F^high^ and Siglec-F^low^ cells from the loss-of-function FLAM library (**Figure 5C**). Following genomic DNA extraction, sgRNA abundances for each sorted population were determined by deep sequencing. To test for statistical enrichment of sgRNAs and genes, we used the modified robust rank algorithm (α-RRA) employed by Model-based Analysis of Genome-wide CRISPR/Cas9 Knockout (MAGeCK). MAGeCK first ranks sgRNAs by effect and then filters low ranking sgRNAs to improve gene significance testing (53). To identify genes that are required for Siglec-F expression we compared the enrichment of sgRNAs in the Siglec-F^low^ population to the Siglec-F^high^ population. The α-RRA analysis identified over 300 genes with a p-value <0.01 and the second ranked gene in this analysis was the target of the screen Siglec-F (**Figure 5E** and **Table S2**). Guide-level analysis showed agreement with all four sgRNAs targeting Siglec-F, with each showing a ten-fold enrichment in the Siglec-F^low^ population (**Figure 5F**). The high ranking of Siglec-F gives high confidence in genome-wide screen results.

Stringent analysis revealed an enrichment of genes with no previously described role in Siglec-F regulation including the TGFβ response regulator USP9x (54) (**Figure 6A**). To identify pathways that were associated among these genes, we filtered the ranked list to include genes that had a fold change of >4 with at least 2 out of 4 sgRNAs and used DAVID analysis to identify pathways and functions that were enriched in our datasets. The top enriched KEGG pathway was the peroxisome, with all core components of peroxisome biogenesis identified as positive regulators of Siglec-F (**Figure 6B**). We further examined other peroxisome-associated (PEX) genes and found that 11 out of 15 PEX genes present in our library were altered greater than two-fold (**Figure 6C**). KEGG pathway analysis identified a significant enrichment in genes associated with lipid metabolism, including glycerophospholipid, inositol phosphate, and ether lipids (**Table S3**). KEGG analysis also found an enrichment of the phagosome pathway which identified several surface receptors associated with phagocytosis in this pathway, including the IgG Fc Receptor 4, the mannose-6-phosphate receptor, and the oxidized low-density lipoprotein receptor, suggesting surface proteins associated with phagocytosis directly modulate the stability of Siglec-F (**Figure 6D**). Examination of enriched UniProt keyword terms using DAVID analysis found a strong enrichment of proteins with oxidoreductase function including several genes associated with cytochromes P450 (CYP), a key regulator of xenobiotic, fatty acid, and hormone metabolism, known to be important in the lung environment (55). Thus, bioinformatic analysis of the top positive regulators of Siglec-F identified pathways that are associated with AM functions.

**Figure 6.**
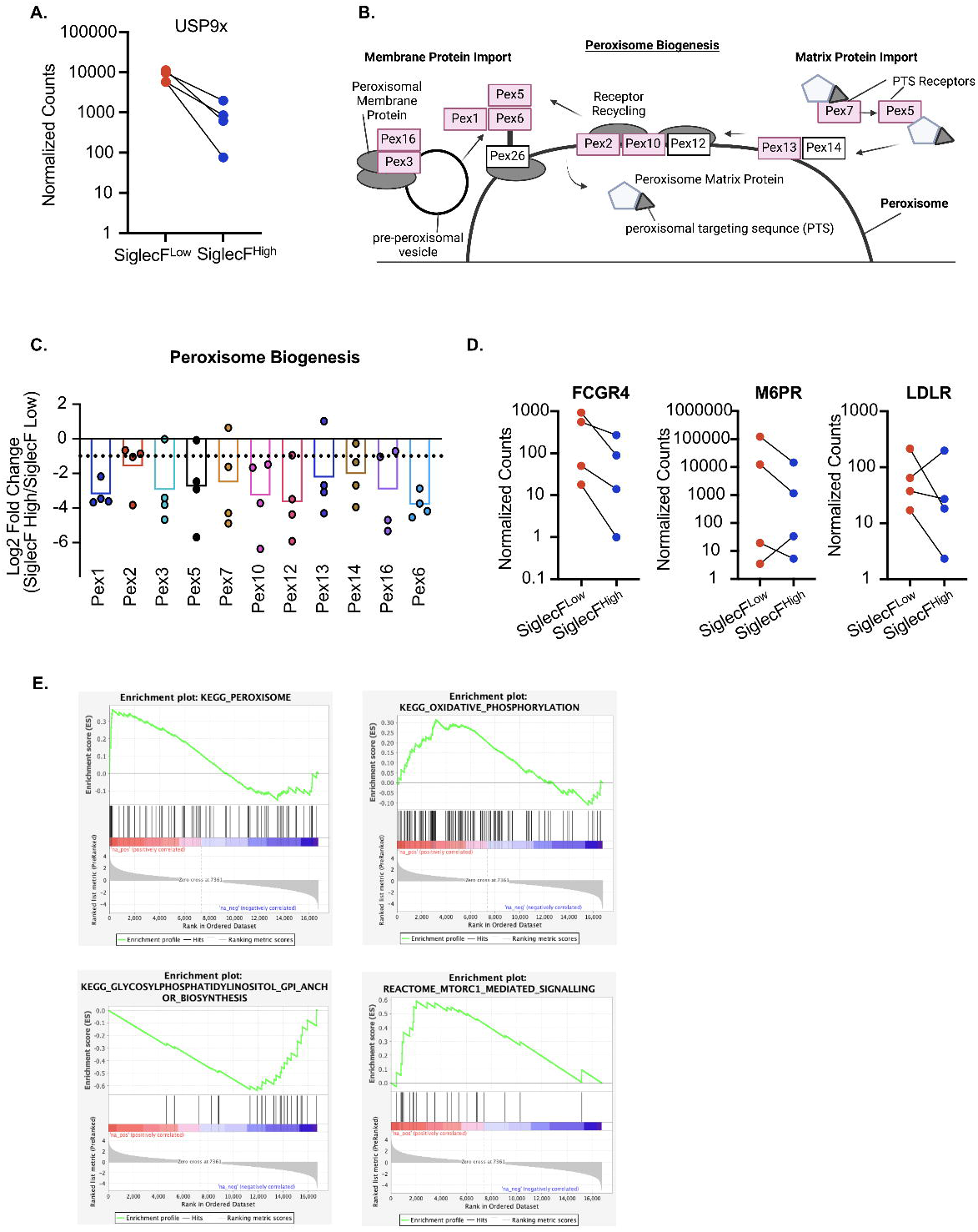
Bioinformatic analysis identifies FLAM metabolic networks as critical regulators of Siglec-F expression. **A)** The TGFβ response regulator USP9x was a significant hit in the screen. Shown are the normalized counts for each of the four sgRNAs targeting USP9x in each sorted population. **B)** Using DAVID analysis, peroxisome biogenesis was identified as the most significantly enriched KEGG pathway. Shown is an adaptation of the KEGG peroxisome biogenesis pathway highlighting the 10 peroxisome regulators identified in the screen in red. **C)** The sgRNA distribution and mean log-fold change for each peroxisome regulator identified in the genetic screen are shown. The dashed line indicates a Log_2_ fold change of −1. **D)** DAVID analysis identified surface proteins associated with phagocytosis. Shown are the normalized counts for each of the four sgRNAs targeting the indicated surface protein from each sorted population. **E)** GSEA was used to identify enriched pathways from the entire forward genetic screen. Shown are four leading-edge analysis plots that are representative of this analysis for a subset of enriched pathways. These pathways including the peroxisome, oxidative phosphorylation, GPI anchor biosynthesis and mTORC1 signaling.

We next used gene set enrichment analysis (GSEA) to identify functional enrichments from the entire ranked screen dataset (**Table S3**). GSEA identified the peroxisome as a top enriched KEGG pathway consistent with the DAVID analysis (**Figure 6E**). This analysis also identified a strong enrichment for oxidative phosphorylation, which is consistent with the key metabolic changes in AMs compared to BMDMs (56), and a significant enrichment for GPI anchor synthesis as negative regulators of Siglec-F surface expression (**Figure 6E**). We also noted that mTORC1 signaling was enriched as a negative regulator, in line with previous reports that mTORC1 is required to maintain AMs in the lungs (57). Taken together, our forward genetic screen not only identified Siglec-F,the screen target, but also identified positive and negative regulators of Siglec-F expression that are associated with known AM-functions as well as novel AM regulators. Thus, FLAMs are a tractable genetic platform that enables the detailed interrogation of AM regulatory functions and mechanisms.

## DISCUSSION

As long-lived resident macrophages in the lungs, AMs have unique phenotypes and functions shaped by the alveolar environment (58). However, experimental limitations hinder our understanding of AM-specific functional mechanisms. Developing *ex vivo* models that recapitulate AM phenotypes would overcome the challenges associated with isolating and maintaining AMs from the lungs of mice. Since AMs are derived from fetal liver monocytes, previous studies tested the culture of fetal liver cells with GM-CSF (59). These culture conditions result in self-replicating AM-like cells, but in our hands, the AM-like phenotype was not stable long-term. While low passage fetal liver cells grown in GM-CSF are useful for some experimental approaches, the instability of the AM phenotype precludes functional genetic studies (60, 61). To stabilize the AM-like phenotype of fetal liver-derived cells we supplemented the growth media with TGFβ, a key cytokine for AM maintenance in the lungs, in a model we term FLAMs (10). Here we showed that FLAMs recapitulate many aspects of AM biology, are stable long-term, and are genetically tractable, making them a useful tool to dissect the regulation of AM maintenance and function.

We demonstrated that that even after one month of culture’ FLAMs efficiently phagocytose cSiO_2_ particles, produce inflammatory cytokines like IL-1α, and die similarly to AMs. Our results are consistent with two recent reports that examined how TGFβ modulates macrophages *ex vivo* (60, 61). These reports showed that TGFβ can induce/maintain AM-like phenotypes *ex vivo* using AMs directly from the lungs of mice or purified cells from the bone-marrow. Other differences in these approaches, including the use of the PPARγ agonist rosiglitazone, make these strategies distinct yet complementary and underscore the key role of TGFβ in AM regulation. The advantage of FLAMs is the low cost, low technology threshold and high yield of cells that can be isolated from any genetically modified mouse. Another key advantage of FLAMs is the potential of the genetic tools developed here. Using targeted gene-editing we showed that directed mutations can be easily generated in FLAMs to probe specific AM functions. Furthermore, we generated a genome-wide knockout library in FLAMs and completed the first forward genetic screen in AM-like cells, highlighting the utility of FLAMs as a genetic platform to probe AM function. Thus, FLAMs recapitulate *ex vivo* AMs even after extended culturing and are suitable for dissecting AM responses and regulation.

How AMs control their functional responses and how this differs from other macrophage populations remains unclear. Here we observed both phenotypic and functional differences among AMs, FLAMs, and BMDMs in line with previous studies (25, 39, 40). While BMDMs express high levels of CD14, AMs and FLAMs express high levels of Siglec-F and MARCO. When cells were exposed to cSiO_2_, we observed differences in cell death kinetics and IL-1 cytokine responses. Though BMDMs were able to engulf cSiO_2_ particles at a rate comparable to AMs and FLAMs, they quickly succumbed to cell death, while AMs and FLAMs remained viable many hours following cSiO_2_ phagocytosis. A delay in cell death may be important for appropriate clearance of particles, potentially allowing the AMs to be transported out of the alveoli before they die (62). AMs and AM-like cells also released significantly more IL-1α and IL-1β than BMDMs in response to cSiO_2_. These data are consistent with studies showing high levels of IL-1α produced by AMs compared to other cells in the lung and other macrophage sub-types (25, 39, 44, 45, 59) and the known role of cSiO_2_ in inducing IL-1β release (49). Our findings indicate that MARCO may be a key player in driving the IL-1 cytokine response in AMs as MARCO-deficient FLAMs showed increased viability and decreased IL-1 production following cSiO_2_ exposure. These results are in line with previous studies implicating MARCO in the uptake of cSiO_2_ and other particles in AMs (50, 63). In the future, FLAMs will be used to dissect the underlying mechanisms of MARCO regulation to understand how MARCO drives distinct inflammatory responses following phagocytosis of cSiO_2_ and other pathogenic cargo. Knocking out the IL-1 receptor also reduced IL-1 cytokine release, which points to a feed-forward mechanism to amplify this inflammatory response in AMs.

In addition to MARCO, AMs express other markers that are used to define AM populations. However, the regulation of these other AM markers, like Siglec-F, remains entirely unknown. Siglec-F is a surface-expressed immunoglobulin protein that binds sialic acid residues on glycolipids and glycoproteins, but its function in AMs is largely unknown. In addition to AMs, Siglec-F is expressed on eosinophils, where it limits inflammation by modulating cell death pathways (64). The only studies examining Siglec-F in AMs demonstrated that Siglec-F does not regulate phagocytic activity (65). Our forward genetic screen in FLAMs defined regulators of Siglec-F surface expression and uncovered hundreds of candidate genes that may contribute to Siglec-F expression. Our results not only identified Siglec-F as the second ranked candidate, but we identified other genes that likely modulate Siglec-F expression or trafficking. These genes include transcription factors like Fos and NFkB2 and surface receptors like M6PR. Our screen candidates are likely to include both direct regulators of Siglec-F expression and indirect regulators that maintain the AM-like state. In support of this prediction, we identified USP9x, a known regulator of TGFβ signaling as a strong positive regulator of Siglec-F expression (54). In addition, we identified the enrichment of functional pathways previously associated with AM function including the peroxisome, lipid metabolism, oxidative phosphorylation and CYP. Given the previous links among PPAR transcription factors, peroxisome biogenesis, and lipid metabolism, our data strongly suggest FLAMs recapitulate the metabolic makeup of AMs which is central to their gene regulation (12, 56, 66). In further relation to the metabolic state of FLAMs, we identified several CYP family members among our top candidates which regulate vitamin A and all-trans retinoic acid, known modulators of AM function (67, 68). Based on these findings, we posit that PPARγ expression drives lipid metabolism to induce the AM-specific transcriptional profile, resulting in Siglec-F expression. Future studies will be centered on testing this model and deeply validating the genetic screen to uncover novel regulatory mechanisms in FLAMs.

FLAMs are a promising model to study AM biology, yet some limitations remain. While FLAMs maintain many AM-like phenotypes long-term, we observed variable expression of the AM marker CD11c over time. This suggests that there are other signals in addition to GM-CSF and TGFβ that are needed to fully recapitulate AM functionality *ex vivo*. The alveolar space is a highly complex microenvironment, with constant crosstalk between AMs and other cells (58). For example, our data show that FLAMs express TGFβ, yet this is not sufficient to maintain AM-like functions and continued *Tgfbr1* and *Tgfbr2* expression. Given TGFβ is known to amplify *Tgfbr1* and *Tgfbr2* this suggests that the TGFβ produced by FLAMs is not biologically active. *in vivo*, latent TGFβ released by AMs is activated by α-V-β-6 integrins expressed on the type II alveolar epithelial cells (AECII), resulting in increased levels of the active protein which can signal in an autocrine manner to maintain the unique phenotype of AMs (69). This feed-forward loop is absent *ex vivo*, which may explain why addition of exogenous active TGFβ prevents the loss of the AM-like phenotype in FLAMs. Other signals provided by AECII cells and others such as lung-resident basophils likely regulate AM maintenance, but these signals are not modeled in our system (70). Combining the genetic tractability of FLAMs with *in vivo* transfer models may enable us in the future to dissect this cross-regulation systematically to further improve FLAMs. Intranasal transfer of TGFβ-cultured AMs was recently shown to repopulate the alveolar space suggesting FLAMs could be similarly instilled, enabling rapid studies to better understand AM- maintenance within the lung environment (27).

In summary, we developed FLAMs, a stable *ex vivo* model that can be used to study lung development, immunology, and toxicology. FLAMs are likely to shed new light on processes unique to AMs, like phagocytosis, efferocytosis and the removal of inhaled particles, by employing targeted or genome-wide genetic approaches. Taken together, the optimization and application of FLAMs provides an exciting, innovative model to thoroughly investigate AM biology.

## Acknowledgements

We would like to acknowledge Dr. Jack Harkema, Dr. Melissa Bates, Dr. Mikhail Givralin, Dr. Alexander Misharin, and the members of the Olive and Pestka lab for helpful discussions and input. We thank the MSU flow cytometry core for their help with instrumentation and analysis and Carol Flegler of the MSU Center of Advanced Microscopy for assistance in scanning electron microscopy.

---

**Figure S1.**
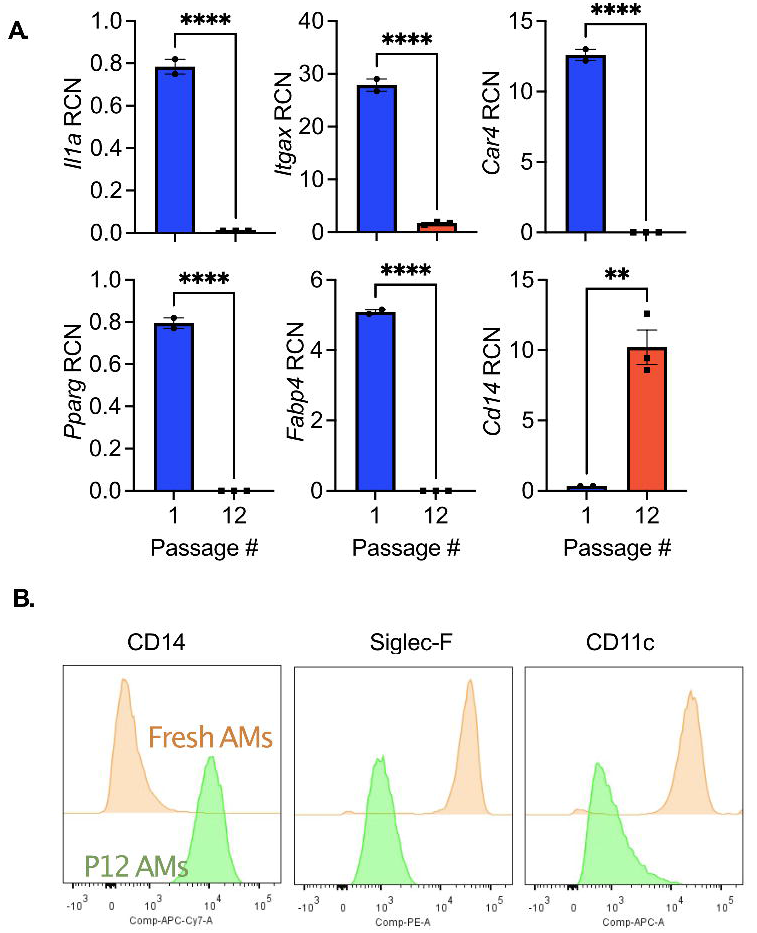
Fetal liver macrophages sorted to enrich for Siglec-F-expressing cells lose the AM-like expression profile. Fetal liver macrophages were cultured for 1 week to allow for development and expression of Siglec-F and CD11c and were then stained and sorted based on Siglec-F expression with over 98% of cells expressing high levels of Siglec-F. Cells were then plated in media containing GM-CSF alone and at the indicated passage cells were lifted and stained for Siglec-F and CD11c Asterisks indicated significant differences between samples (\*\*\**p*<0.001) as determined by a student’s T-test. Results are representative of two independent experiments.

**Figure S2.**
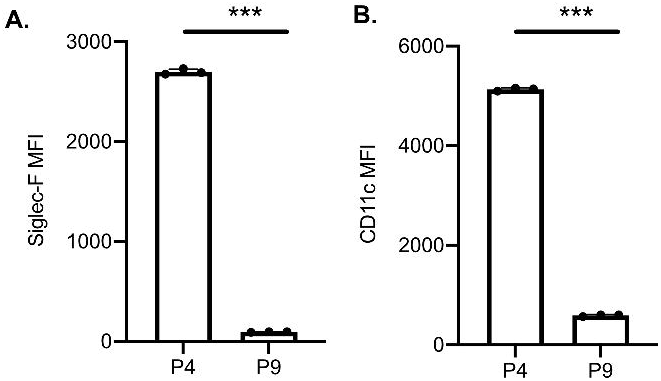
Long term *ex vivo* culture of AMs leads to loss of AM-defining genes and surface markers. Alveolar macrophages (AM) were isolated and cultured in RPMI with 20 ng/mL GM-CSF for approximately 2 months (12 passages). Gene expression **(A)** and cell surface markers **(B)** were compared between cultured AMs and freshly isolated AMs. Asterisks indicate significant differences (\*\*\*\**p*<0.0001, \*\**p*<0.01) between samples.

**Figure S3.**
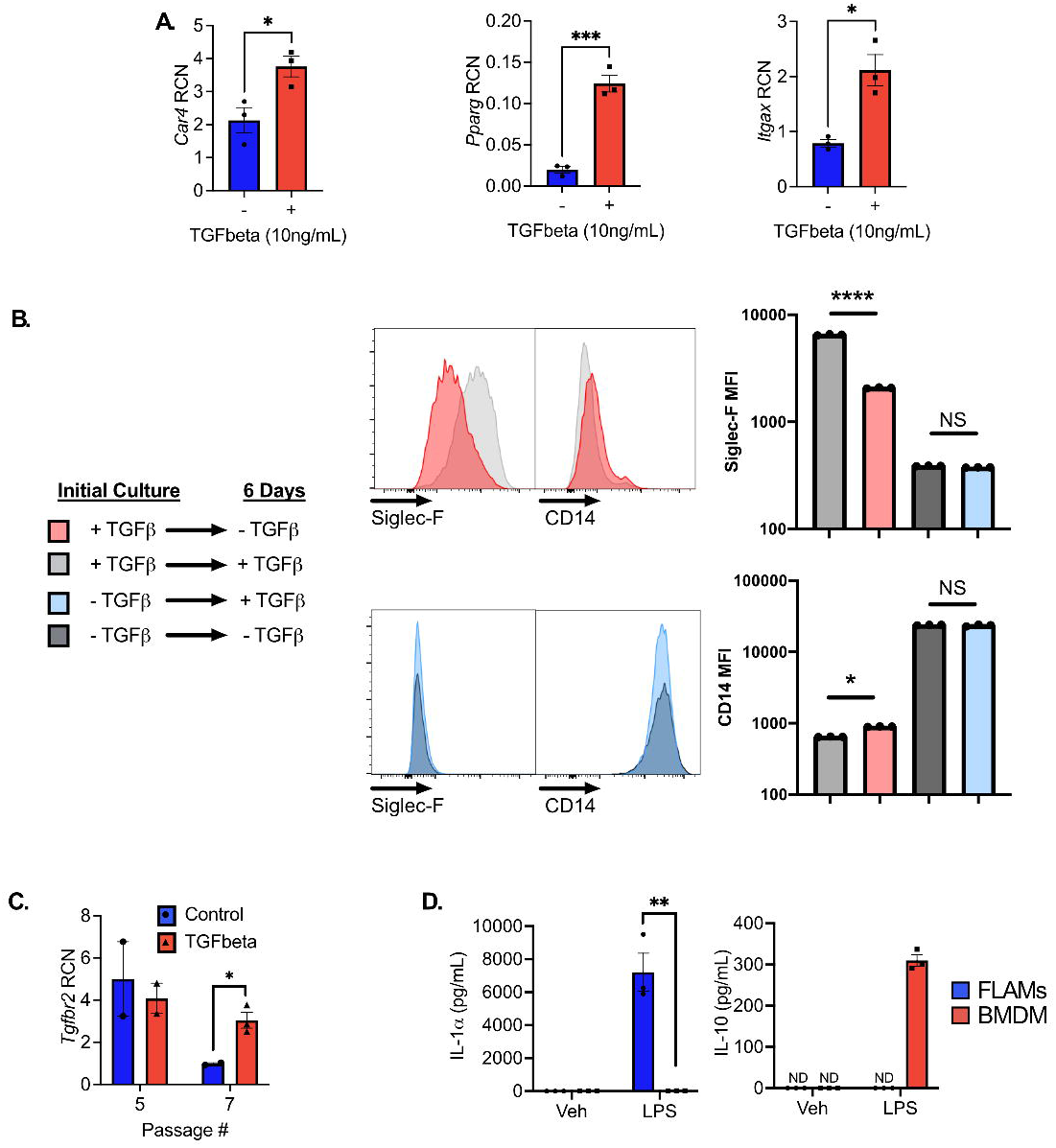
TGFβ induces and maintains AM-like state in early-passage but not older fetal liver macrophages. **A)** Freshly isolated fetal liver cells were incubated with 10 ng/mL TGFβ for 24 hours prior collecting RNA for qPCR analysis of *Car4, Itgax* and *Pparg* gene expression. **B)** Fetal liver macrophages that were low passage (<4) and kept in TGFβ and those that were high-passage (>15) and cultured without TGFβ were cultured for 6 days with and without TGFβ. On day 6, cells were lifted and surface expression of Siglec-F and CD14 was assessed by flow cytometry. **C**) Expression of *Tgfbr2* in fetal liver macrophages cultured with and without TGF-β was assessed at P5 and P7. Asterisks indicate significant differences (****p<0.0001, ***p<0.001, *p<0.05) between samples, as determined by Student’s t-test. **(D)** Early fetal liver-derived cells and bone marrow-derived macrophages (BMDMs) were treated with 100 ng/mL LPS for 24 hours and cell-free supernatant was analyzed for release of IL-1α and IL-10 by ELISA. Asterisks indicate significant differences (**p<0.01) between fetal liver macrophages cells and BMDMs. ND=not detected.

**Figure S5.**
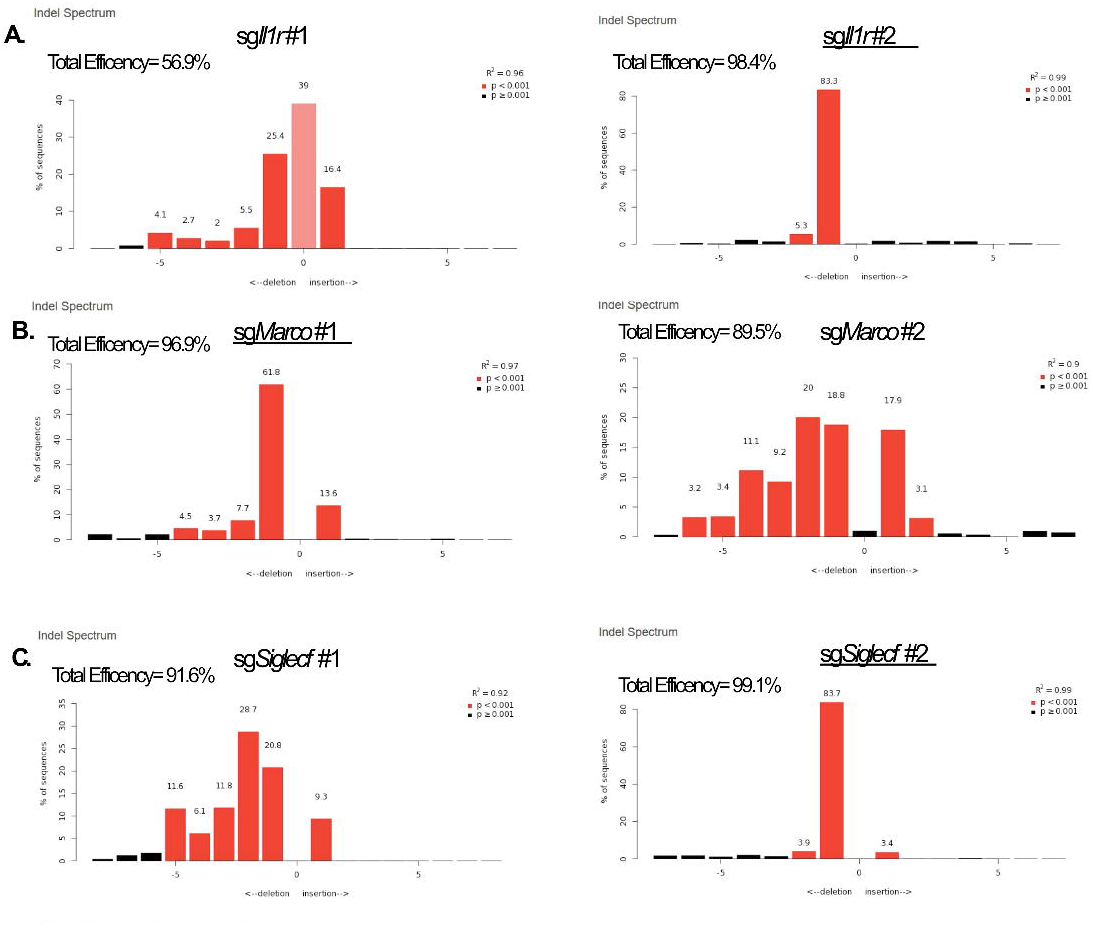
Quantification of editing efficiency in FLAMs targeted with CRISPR-Cas9. Cas9^+^ FLAMs were transduced with lentivirus carrying sgRNAs targeting the indicated gene. Following selection, genomic DNA was isolated from control and targeted cells and PCR was used to amplify the region encoding each targeted gene. TIDE analysis was used to quantify the editing efficiency of the indels in each cell line using trace plots following Sanger sequencing. Shown is the TIDE analysis profile indicating the percent editing efficiency for each sgRNA targeting A) *Il1r1* B) *Marco* and C) *Siglecf*. Red bars indicate the presence of indels of the indicated size. Knockout cells used the remainder of the manuscript in Figures 5 and 6 are underlined.

**Table S1. Quantification of the coverage in the genome-wide knockout library. (Sheet1)** Statistics for the CRISPR screens and zero count sgRNAs are quantified. (**Sheet2**) Normalized counts for each sgRNA from two biological replicates of the genome-wide knockout library are listed.

**Table S2. Gene and sgRNA analysis for the Siglec-F genome-wide screen. (Sheet1)** The gene summary output from MAGeCK RRA analysis is shown based on SiglecF^high^/SiglecF^low^. (Sheet2) The sgRNA summary output from MAGeCK RRA analysis is shown for each sgRNA in the library.

**Table S3. Bioinformatic analysis of Siglec-F genome-wide screen.** Each labeled sheet indicates the type of analysis (DAVID or GSEA) and the lists the pathways or genes that are associated with each analysis.

